# Successive responses of three coral holobiont components (coral hosts, symbiotic algae, and bacteria) to daily temperature fluctuations

**DOI:** 10.1101/2023.09.30.560297

**Authors:** Yunli Eric Hsieh, Chih-Ying Lu, Po-Yu Liu, Jia-Min Kao, Sung-Yin Yang, Chien-Yi Wu, Jing-Wen Michelle Wong, Shinya Shikina, Tung-Yung Fan, Shan-Hua Yang

**Affiliations:** Systems Biology and Mathematical Modelling, Max Planck Institute of Molecular Plant Physiology, Potsdam, Germany; Bioinformatics, Institute of Biochemistry and Biology, University of Potsdam, Potsdam, Germany; School of BioSciences, The University of Melbourne, Parkville, Australia; Molecular and Biological Agricultural Sciences Program, Taiwan International Graduate Program, National Chung Hsing University and Academia Sinica, Taipei, Taiwan; Graduate Institute of Biotechnology, National Chung Hsing University, Taichung, Taiwan; Biodiversity Research Center, Academia Sinica, Taipei 11529, Taiwan; Department of Post-Baccalaureate Medicine, National Sun Yat-sen University, Kaohsiung, Taiwan; Institute of Fisheries Science, National Taiwan University, Taipei, Taiwan; Department of Aquatic Bioscience, National Chiayi University, Chiayi, Taiwan; Institute of Marine Environment and Ecology, National Taiwan Ocean University, Keelung, Taiwan; National Museum of Marine Biology and Aquarium, Pingtung, 944, Taiwan

**Keywords:** Reef coral, *Stylophora pistillata*, *Pocillopora acuta*, daily temperature fluctuations, microbiome, successive changes

## Abstract

Coral reef ecosystems support over a quarter of the world’s marine life and play important ecological and economic roles. However, the increasingly severe weather events associated with ocean warming and climate change are believed to be rapidly altering the functions of coral reefs and their ecosystems. Corals and their associated microbiota form a “holobiont,” which includes symbiotic algae and other associated microbiota dominated by bacteria. These microbiota have a direct relationship with the health of the coral host. Their composition is influenced by various environmental factors, such as increasing sea water temperatures. Previous studies of the effects of temperature changes on coral physiology and associated bacterial communities have been conducted based on stable water temperatures set by mean temperatures, or by slowly increasing/decreasing temperatures. However, the daily temperature fluctuations that corals experience in nature are not stable. Rather, there may be significant differences of up to 6°C in a single day. The current understanding of the effects of large daily temperature fluctuations on coral and associated bacterial community dynamics is limited. Hence, in this study, we conducted a four-week tank experiment using different large daily temperature fluctuations accompanied by continuous warming conditions to investigate the effects on two common reef-building corals, *Stylophora pistillata* and *Pocillopora acuta*, in Taiwan. During the experiment, the activity of coral host catalase was measured, the photosynthetic ability of symbiotic algae was recorded, and the variation in bacterial communities was analyzed using the V6-V8 region of 16S rDNA. According to the results, different parts of the holobionts of different coral species exhibited varying response rates to the continuous warming conditions and diurnal temperature fluctuations. Additionally, it was found that diurnal temperature fluctuations may mitigate the heat stress on the host and reduce the changes in bacterial response to warming. Furthermore, the holobionts of different coral species may adopt different adaptation and survival strategies in response to diurnal temperature fluctuations and warming. Finally, based on the response of these two coral species under the conditions of diurnal temperature fluctuations and continuous warming, *Acinetobacter* and *Rhodobacteraceae* were identified as potential indicator coral-associated bacteria. This is the first study to investigate the tripartite dynamic response of coral, symbiotic algae and bacteria to daily temperature fluctuations.

## 1. Introduction

Coral reef ecosystems harbor 25% of the world’s marine organisms and play essential ecological and economic roles (National Oceanic and Atmospheric Administration, NOAA). Global climate change has led to seawater surface temperature (SST) anomalies, which threaten coral reef ecosystems and have already caused several mass coral bleaching events (Berkelmans et al., 2004; Costanza et al., 2014; Hughes et al., 2017). The bleached corals are susceptible to disease, and prolonged bleaching can cause coral mortality (Berkelmans et al., 2004; Hughes et al., 2017; Hughes et al., 2018) The estimated cost in damages to this ecosystem on which human societies depend is almost US $36 billion per year (Spalding et al., 2017).

All biomes are fundamentally dependent on their microbial constituents (Azam and Worden, 2004). Diverse microorganisms, mainly the symbiotic algae, Symbiodiniaceae, and bacteria, are harbored by coral and form an integrated holobiont. These microorganisms have diverse interactions with their host and maintain coral holobiont functions, such as nutrient acquisition and health regulation (Rosenberg et al., 2007). It is well known that symbiont algae are the main carbon producers in coral. In addition, coral-associated bacteria are the most dominant bacteria in corals, and they support other essential nutrient cycling pathways (Lema et al., 2012; Yang et al., 2016). The composition of the coral holobiont is influenced by various micro and macro environmental factors, such as increasing SST, which has caused the dysbiosis of Symbiodiniaceae, bacteria, and coral hosts and led to bleaching and mortality.

Numerous studies have relied upon average annual or summer threshold temperatures to produce predictions and scenarios about the effect of thermal stress on corals and coral bleaching (Hughes et al., 2017), despite shallow corals in situ normally experiencing highly dynamic diurnal temperatures in intertidal zones (Easterling et al., 2000; IPCC, 2014). It has been suggested that corals’ physiological tolerance and performance in response to thermal stress could be influenced by historical diurnal temperature fluctuation (Castillo et al., 2005; Carilli et al., 2012; Palumbi et al., 2014). Safaie et al. (2018) presume that increasing the short-term temperature range could reduce the risk of coral bleaching, based on examples of 20 environmental variables and 81 bleaching events in 5 major reef regions globally. However, the precise temperature range within which corals can acclimatize and develop resilience to bleaching is still unclear.

Daily temperature fluctuations have the potential to affect, or even disrupt, the scale and direction of biotic interactions, including community dynamics (Gilman et al., 2010) and ecosystem functions (Traill et al., 2010). Accurately predicting interactions within holobionts and among species in an ecosystem under a dynamic environment requires an in-depth understanding of the species-specific responses to fluctuating environmental conditions. To date, the majority of studies have examined the impacts of environmental changes on coral hosts, algal symbionts, and bacteria separately. Only a few studies have investigated successive reactions within a holobiont during stable increasing temperature conditions (Li et al., 2021). Hence, it remains uncertain which part of the holobiont will be most affected by and sensitive to varying degrees of diurnal temperature fluctuations.

To investigate the response of host, symbiotic algae, and bacterial communities to a variety of dynamic temperature conditions, we conducted a four-week tank experiment using different large daily temperature fluctuations accompanied by continuous warming conditions on two common shallow reef-building corals, *Stylophora pistillata* and *Pocillopora acuta*, in Taiwan. These two coral species are commonly used for coral physiology experiments and have been known to have different thermo-tolerant performances. Notably, *Pocillopora acuta* is found to acclimatize to heat-stress better than *S. pistillata* at Outlet reef in southern Taiwan (Keshavmurthy et al., 2014). By analyzing the catalysis activity of hosts, examining the photosynthetic activity of symbiotic algae, and investigating coral-associated bacterial dynamics using the V6-V8 region of 16S rDNA, we hoped to understand the successive changes of hosts, symbiotic algae, and bacteria to large daily temperature fluctuations. Furthermore, we hoped to identify bacteria that are sensitive to daily temperature fluctuation and thus could serve as indicator genera or species.

## 2. Materials and methods

In this experiment, we treated coral with two daily temperature ranges that can cause coral bleaching: 26°C±5°C and 26°C±7°C. To explore the coral physiological condition during the treatments, we measured the Catalase (CAT) level and photochemical efficiency of the coral tissue. The bacteria compositions in coral tissue were further analyzed throughout the entire experiment period.

### 2.1 Sample collection and incubation experiment

*S. pistillata* were collected from Bitoujiao Park, northern Taiwan (25.126263 N, 121.914244 E) in January 2020, and *P. acuta* were collected from Outlet reef, southern Taiwan (21.930722 N, 120.745250 E) in July 2020. Each colony was cut into 12 fragments about 3 cm long and were then glued on a tile base and evenly distributed into 3 indoor, closed system seawater tanks for acclimation at 26 °C for 1 week. After acclimation, temperatures were set as (groups A and D) 26-29°C, (groups B and E) 26±5°C-29±5°C, (groups C and F) 26±7°C-29±7°C, controlled by the APEX system (Apex System, Neptune, USA) with aquarium heating rod (ADP-350W, Taiwan) and aquarium chiller (IPO-300, Taiwan), and recorded by HOBO temperature loggers (HOBO Pendant 13 Temp/Light, Onset, USA). Illumination was controlled using LED lights with a daily light intensity of 150 μmol/m^2^/s over a 12-hour light and 12-hour dark cycle (Optimus Reef Nano, MMC PLANNING Co. Ltd, Japan). Thewater in the tanks contained a mixture of artificial seawater (Coral Reef Pro Salt, COVE, Netherlands) and reverse osmosis water (Milli-Q Integral, Merck, Germany) to make the salinity fall to 32-33‰. The water was changed twice a week by exchanging 10% of the tank water with artificial seawater (Coral Reef Pro Salt, COVE, Netherlands). Water pH was monitored using a probe (Apex Systems, pH probe, Neptune, USA). Calcium, magnesium, and carbonate hardness were also measured routinely using Salifert test kits (Calcium Profi Test Kit; Magnesium Profi Test Kit; Carbonate Hardness & Alkalinity Test Kit, Salifert, Nederland). If any of these elements were found to be insufficient, the tanks were supplemented with calcium chloride, magnesium dichloride, or sodium bicarbonate. Newly hatched brine shrimp were used to feed the coral nubbins once per week and maintain the coral nutrition requirement.

### 2.2 Record of appearance and measurement of photochemical efficiency

The treatments were conducted over four weeks. On the last day of each week, the corals were taken up for physiological and microbiome analysis. First, the survival status of corals was checked according to whether their tentacles were stretching out and whether the tissue was festering. Second, the photosynthesis index of the corals was assessed by measuring the maximum yield (Fv/Fm) with a Diving PAM (DIVING-PAM Underwater Fluorometer, Heinz Walz GmbH, Germany), after the coral fragments adapted to the dark for 30 minutes. Third, the coral bleaching condition was checked by comparing the coral color against the coral health chart (Coral Health Chart, Coral Watch, The University of Queensland, Australia) and then photographed. Finally, the coral fragments were rinsed with sterilized artificial seawater, and 2-cm-long branches were cut down with bone scissors, soaked in a 50 ml centrifuge tube with 99% ethanol, and then stored in a -20°C refrigerator for DNA extraction.

### 2.3 Catalase (CAT) activity measurement

We followed Krueger et al.’s (2015) method for the collection of coral tissue homogenate. The method, in brief, went as follows: an airbrush loaded with an ice-cold lysis buffer (50 mM phosphate, 0.1 mM EDTA, 10% [v/v] glycerol, pH 7.0) was used to wash the tissue out of the skeleton. The collected tissue homogenate was kept on ice and then centrifuged at 2000 g for 5 min at 4°C. The supernatant was collected and centrifuged again at 16000 g for 5 min at 4°C. The resulting supernatant was aliquoted, snap-frozen with liquid nitrogen, and stored at -80°C.

The host CAT activity was determined by measuring the depletion of H_2_O_2_ at 240 nm with a spectrophotometer (U-3900, Hitachi, Japan). A blank was prepared by mixing 740 μl of potassium phosphate buffer (50 mM, pH 7.0, 0.1 mM EDTA) with 10 μl of the sample in a quartz cuvette. The reaction began with the addition of H_2_O_2_ to a final concentration of 20 mM and was monitored for 3 minutes at room temperature. Triplicate measurements were made for each sample, and the CAT activity was calculated with an extinction coefficient of 43.6 M^-1^ cm^-1^ (Beers and Sizer, 1952). The protein content of the samples was quantified using the Pierce™ 660 nm protein assay kit (Thermo Fisher, US) with BSA as the standard. To test the difference between treatments, two-way mixed ANOVA was conducted using R.

### 2.4 Coral sampling, DNA extraction and PCR

Coral tissues were collected and sprayed with an airbrush full of 10X TE buffer (10 mM Tris–HCl, pH 8.5, 1 mM EDTA, pH 8.0). After washing the tissue pellets with 10X TE buffer and discarding the supernatant, the tissue pellets were transferred to PowerBead Tubes in the DNeasy PowerSoil Kit (Qiagen, Germany) for total DNA extraction. A PCR was performed using two bacterial universal primers, 968F (5’-AACGCGAAGAACCTTAC-3’) and 1391R (5’-ACGGGCGGTGWGTRC-3’) (Yarza et al., 2014), specifically designed for targeting the bacterial V6-V8 hypervariable regions of the 16S ribosomal rDNA. The PCR protocol consisted of 30 cycles with an initial step of 94°C for 5 min, 94°C for 30 s, 52°C for 20 s, 72°C for 45 s, and finally 72°C for 10 min. Each PCR product was tagged using a DNA tagging PCR method (Chen et al., 2011), then sequenced using the Illumina Miseq 300 bp paired end configuration.

### 2.5 Amplicon Sequence Analysis with KTU Re-Clustering

The 16S rDNA amplicon sequences were processed using the Quantitative Insights Into Microbial Ecology 2 (QIIME 2) pipeline (version 2019.10) (Bolyen et al., 2019). All raw reads were first demultiplexed by cutadapt (version 1.15) (Martin, 2011), then the demultiplexed sequences were denoised by the DADA2 algorithm with quality filtering (by truncating both ends of the reads to 235 bp) and chimera removal (Callahan et al., 2016). The qualified amplicon sequence variants (ASVs) were then assigned taxonomy using the classifier-consensus-vsearch plugin (Bokulich et al., 2018) and the SILVA 128 NR99 database (Quast et al., 2013; Yilmaz et al., 2014). The ASV sequences were then re-clustered using the “K-mer-based taxonomic (KTU) clustering algorithm” (Liu et al., 2022) to refine the sparseness of the ASV abundance table. The unassigned taxon, chloroplast, and mitochondria reads were manually removed from the KTU table.

### 2.6 Statistical Analysis

The bacterial community analyses were conducted and visualized using the vegan (Oksanen et al., 2015) and MARco R-packages (Liu, 2021). A Kruskal-Wallis test or Mann–Whitney U test was used for all statistical analyses of group comparisons under a criterion of type I error α=0.05, and a Dunn’s test was used for post-hoc comparisons in R software (R Core Team, 2015). Dissimilarities among microbial communities were measured by Aitchison distance using a principal coordinates analysis (PCoA), and heterogeneity was tested using ADONIS (permutational multivariate analysis of variance using distance matrices). Alpha diversity indices were estimated by richness, Shannon’s index, Simpson’s index and Chao1 index. An indicator species analysis was done with a chi-square test (Niemi et al., 1997); the indicator species were then identified by Pearson residuals > 5. The differential abundance taxa or indicator species among experimental conditions were visualized with a heatmap using the pheatmap R-package.

## 3. Results

The color of *S. pistillata* was stable throughout the experiment period for both the control and the ±5°C treatment. Although the survival rate of control, ±5°C treatment and ±7°C treatment are 100%, bleaching started to occur in the third week for the ±7°C treatment group, and complete bleaching was observed in the last week (Fig. 1). The changing color of *S. pistillata* aligned with the drop in its photosynthetic activity (Fig. 2a, 2b). Initially, the average Fv/Fm value was around 0.7 for the ±7°C treatment group, but it decreased to 0.5in the third week and to 0 in the last week. This decline in photosynthetic activity may have been the synergetic effects of the increased and fluctuating temperature. Notably, the physical response of Symbiodiniaceae in *S. pistillata* appeared only from the third week, suggesting that it could depend on the amplitude of the daily temperature fluctuation.

**Fig. 1.**
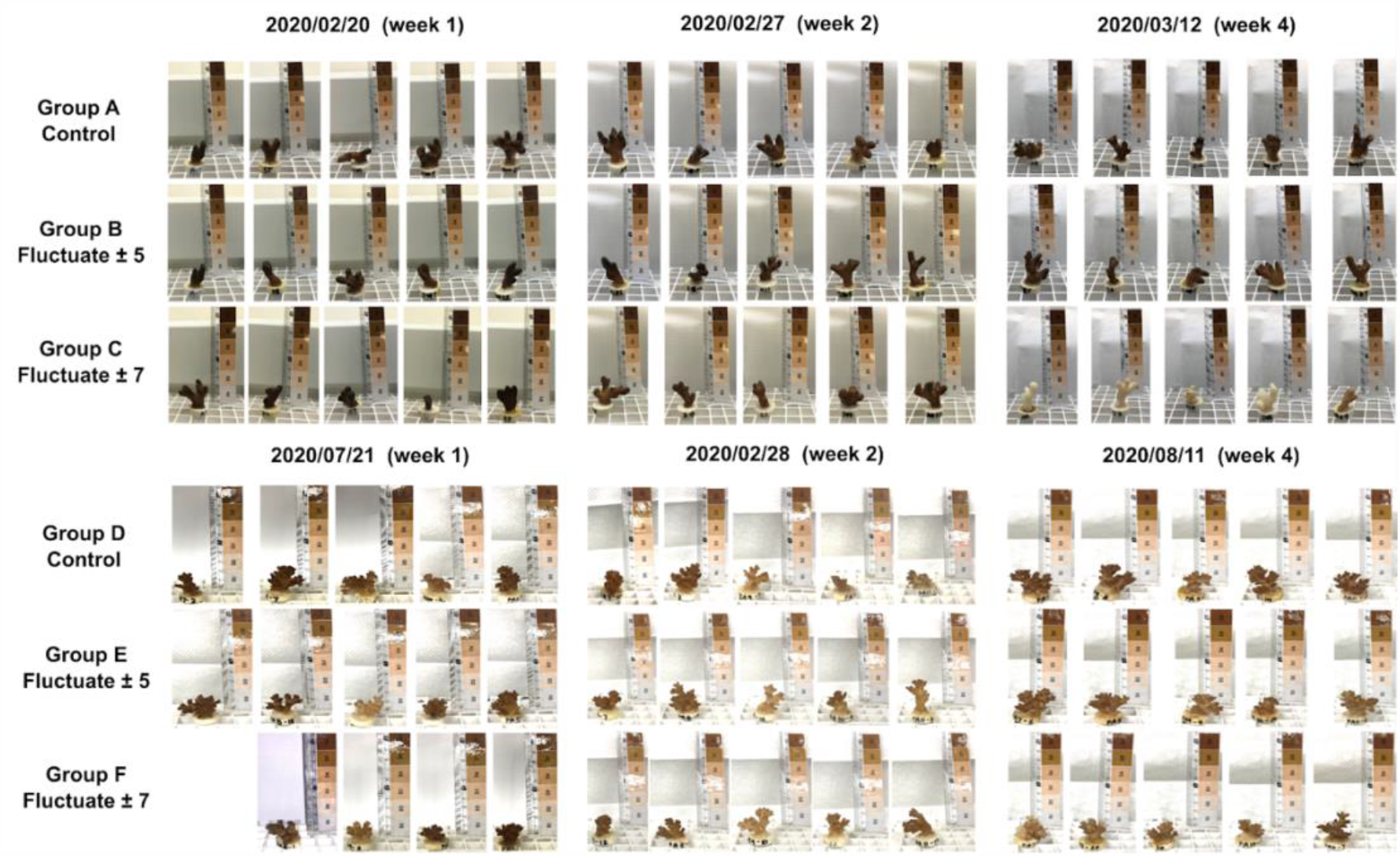
Coral bleaching status during the experiment. The color of corals was checked weekly against the coral health chart. *S. pistillata* is in groups A, B, and C; and *P. acuta* is in groups D, E, and F.

**Fig. 2.**
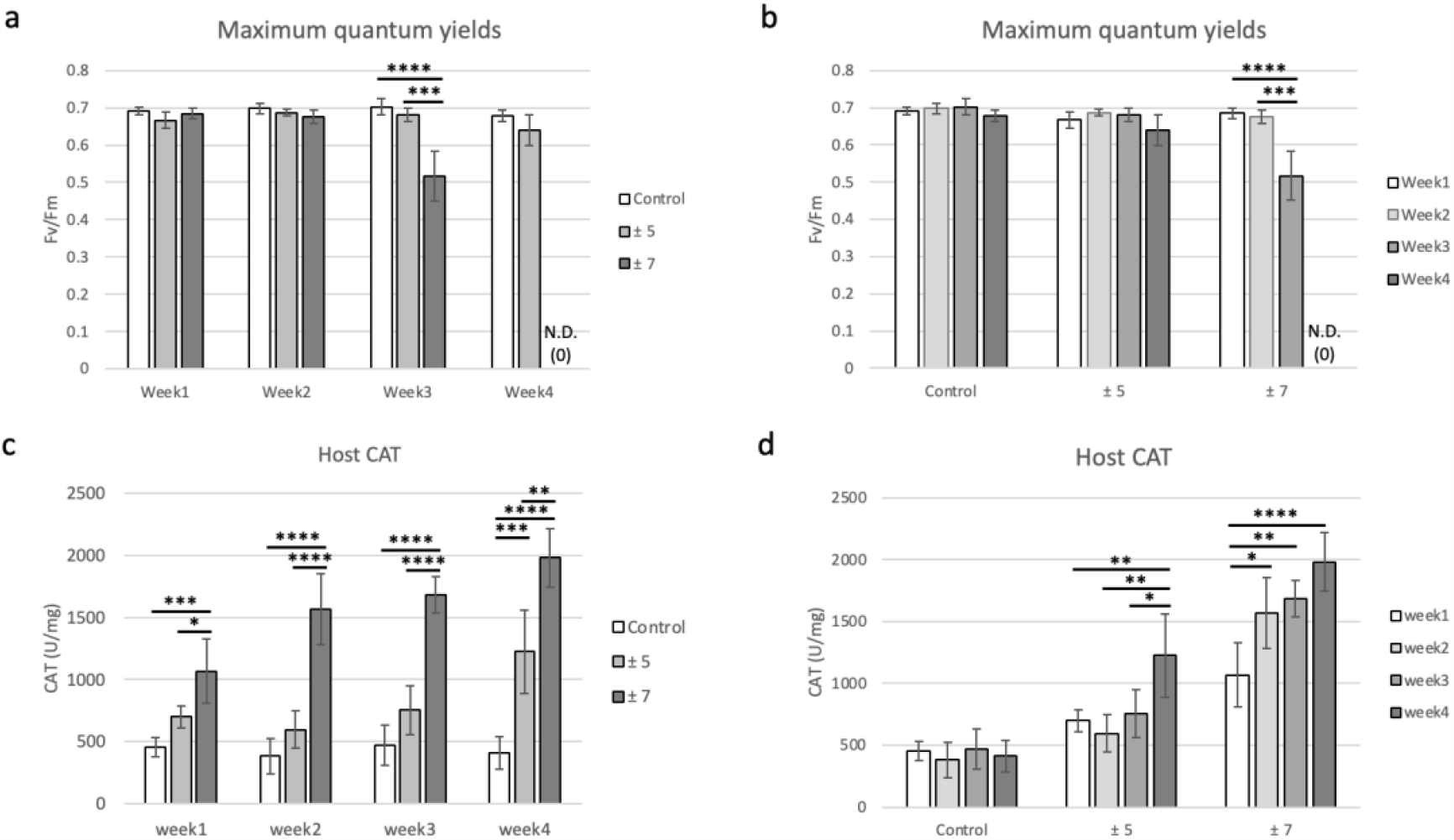
PAM and CAT of *S. pistillata*. a and c show the Fv/Fm comparison and CAT results of the treatment groups across each week. b and d show the Fv/Fm comparison and CAT results of each treatment group throughout the entire duration.

The effect of daily temperature fluctuation on *S. pistillata* can be observed from the CAT results (Fig. 2c, 2d). In the first week, the host CAT activity in both the ±5°C group and ±7°C group was higher than in the control group (Fig. 2c). While the CAT was around 500 U/mg for the control group throughout the experiment period, it was around 700 U/mg in the ±5°C group for the third week and then increased to around 1250 U/mg in the fourth week. In the ±7°C group, the CAT response was double the control group’s in the first week, it increased to 1500 U/mg in week two and week three, and then finally reached around 2000 U/mg, which was significantly higher than the control (Fig. 2c, 2d). This indicates that the host may have been responsive to both the ±5°C and ±7°C treatments from the beginning, but in different magnitudes.

On the contrary, the control and all treatments of *P. acuta* showed no sign of visual bleaching during the four-week period (Fig. 1), nor did they show a decrease in photosynthetic activity (Fig. 3a, 3b). The physiological response of Symbiodiniaceae on warming and daily temperature fluctuation is not significant in *P. acuta*.

**Fig. 3.**
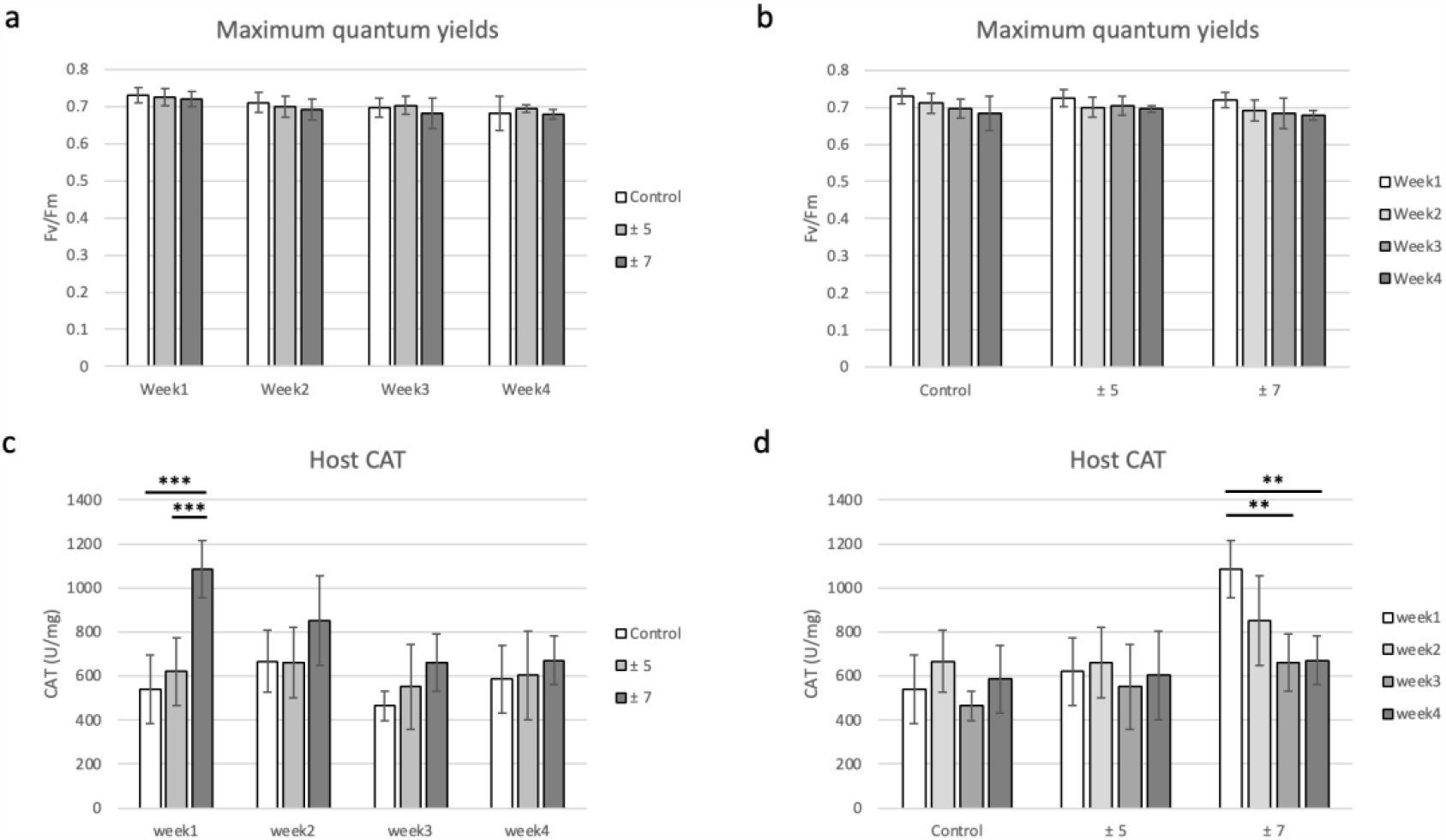
PAM and CAT of *P. acuta*. a and c show the Fv/Fm comparison and CAT results of the treatment groups across each week. b and d show the Fv/Fm comparison and CAT results of each treatment group throughout the entire duration.

The *P. acuta* host was affected in similar ways by the control and the ±5°C group, as CAT levels remained stable around 500 to 600 U/mg throughout the entire four-week period (Fig. 3c, 3d). The temperature fluctuation of the ±7°C group stimulated the highest CAT level of 1100 U/mg at the initial week, which then gradually decreased to a similar level to the control and ±5°C group after the second week (Fig. 3c, 3d). Thus, it appears that the host could have been initially affected by the daily temperature fluctuation but then subsequently acclimated to it. Despite the higher level of warming, the performance of the ±7°C group was not significantly different from the control group, but it still reacted dramatically throughout the experiment period. This suggests that the impact of warming on *P. acuta* is not significant but that the host is more affected by the daily temperature fluctuation.

The two coral species harbor distinct coral microbiome communities (Fig. 4), demonstrating host specificity. However, the microbial communities within these two coral species exhibited variations, with greater temperature fluctuation leading to a convergence in the beta diversity of the microbial communities. The magnitude of this convergence varied depending on the host species.

**Fig. 4.**
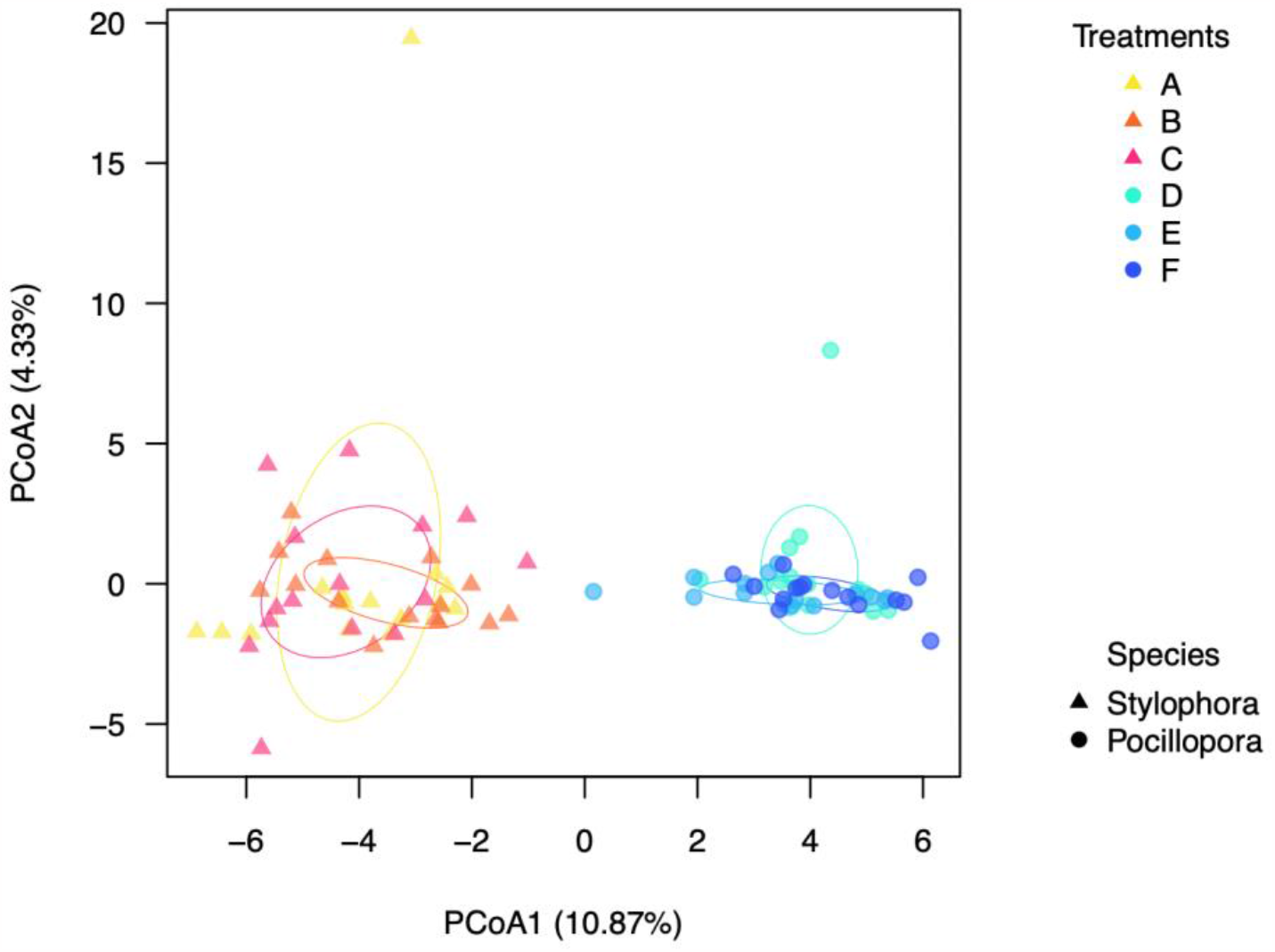
Beta diversity of bacterial KTUs from different treatments in *S. pistillata* and *P. acuta*. Triangles indicate bacterial KTUs in *S. pistillata*, and circles indicate bacterial KTUs in *P. acuta*. Aitchison distance was used for Principal Coordinates Analysis (PcoA). Temperatures were set as (groups A and D) 26-29°C, (groups B and E) 26±5°C-29±5°C, (groups C and F) 26±7°C-29±7°C.

In the case of the *S. pistillata* microbiome, the effect of the ±5°C daily fluctuation treatments was, overall, not significantly different from that of the control group (Fig. 5a), though they did differ at points. In weeks 1 and 2, the ±5°C group showed no significant difference from the control (PERMANOVA R=0.11, p=0.51 in week1; R=0.12, p=0.26 in week2) (Fig. S2a & c), but in week 4, the ±5°C group did significantly differ from the control (PERMANOVA R=0.2, p=0.017) (Fig. S2e). Additionally, although there was no significant difference between the ±7°C group and the control (PERMANOVA R=0.15, p=0.077 in the week) (Fig. S2b), the microbiome of the ±7°C group started to be different from that of the control in the first week (Fig. S1a). The microbiome further began to change due to warming from the second week onwards (PERMANOVA R=0.17, p=0.009 in the week) and neither of the results from week 2 or week 4 overlapped with those of week 1 (Fig. 5c), though these were not significant. Therefore, it can be concluded that the effect of warming on the microbiome will be different depending on the degree of daily temperature fluctuation. If the daily temperature fluctuation is small, the response to warming may be slow, similar to that of the symbiotic algae. If the daily temperature fluctuation is large, the response to warming will be as fast as that of the host.

**Fig. 5.**
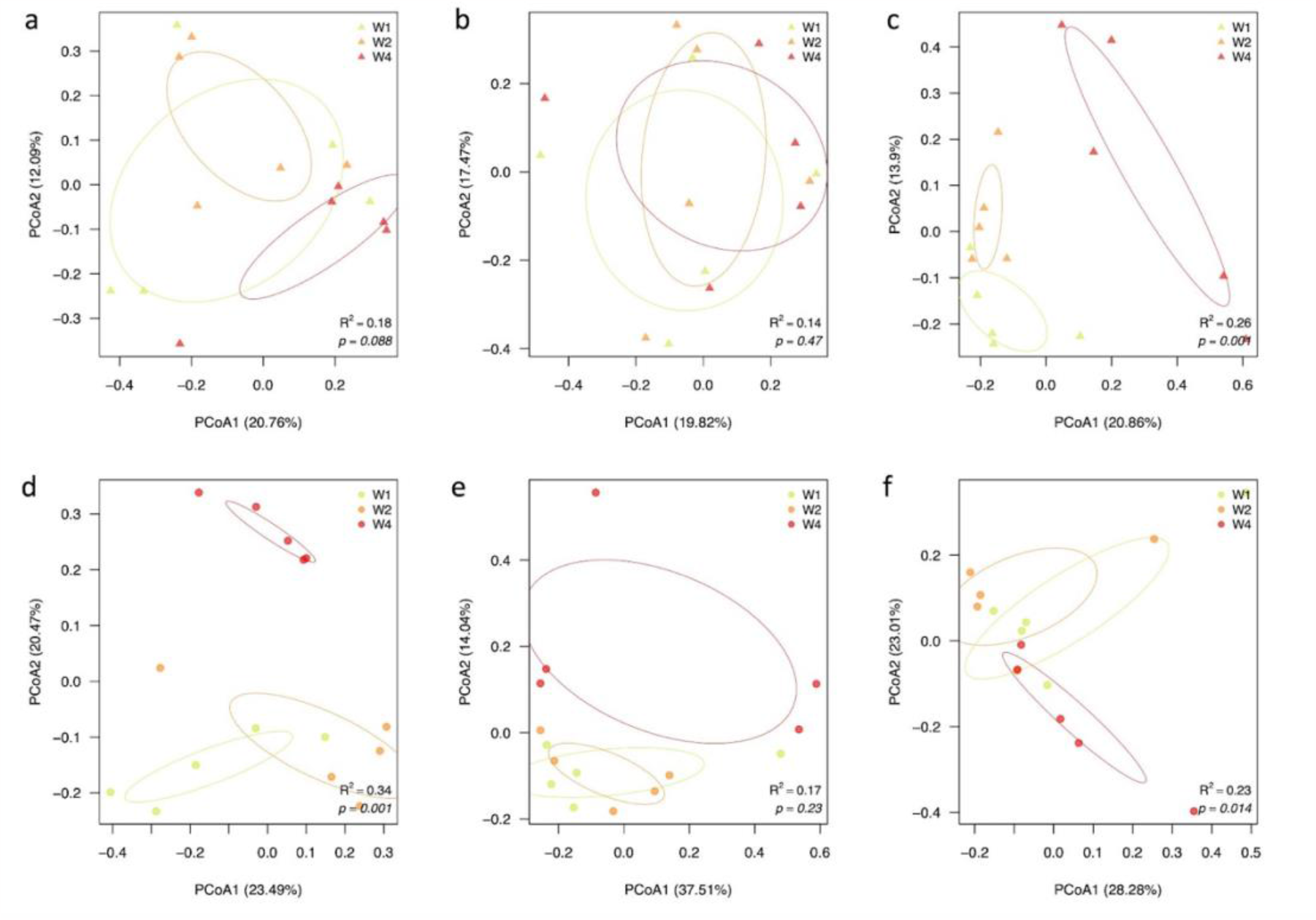
Bacterial dynamics of different temperature treatments. Panels a-c show *S. pistillata* associated bacterial results in groups A (26-29°C), B (26±5°C-29±5°C), and – C (26±7°C-29±7°C), respectively. Panels d-f show *P. acuta* associated bacterial results in groups D (26°C-29°C), E (26±5°C-29±5°C), and F (26±7°C-29±7°C), respectively.

We can therefore likely conclude that in *S. pistillata* under this treatment, the host is the most sensitive, followed by the microbiome, then the symbiotic algae (Table 1).

**Table 1.**
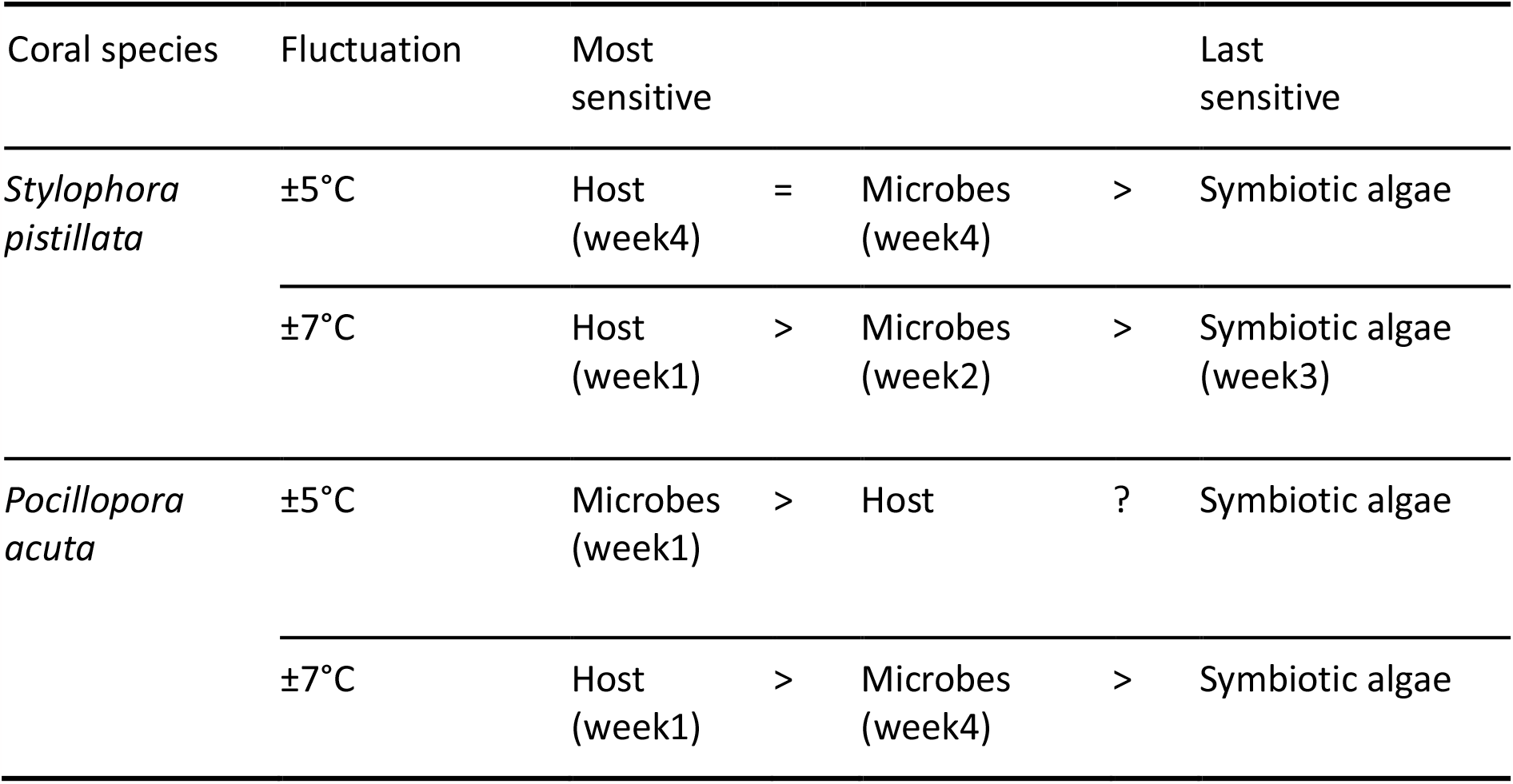
Comparison of coral holobiont components reacting to the increasing temperature with daily fluctuation.

We also investigated the effect of daytime temperature fluctuation on the *P. acuta* microbial community. The microbial communities in the groups subjected to daytime temperature fluctuations showed differences from that of the control group (Fig. S1b). In the first week, the ±5°C group showed significant difference from the control (PERMANOVA R=0.24, p=0.009) (Fig. S3a), but the ±7°C group showed no significant difference to the control (PERMANOVA R=0.16, p=0.082) (Fig. S3b).

As the temperature increased, there were significant differences in the microbial community of the control group at the three measured time points. However, the overall differences in the microbial community in the ±5°C group were not significant, although there was little overlap in the fourth week compared to the first two weeks (Fig. 5b). In the ±7°C group, although the microbial community changed significantly over time, some of the microbial community in the fourth week was still similar to that of the first and second weeks (Fig. 5b). These results suggest that the effect of warming on a microbial community changes depending on the degree of daytime temperature fluctuation. In addition, the presence of daytime temperature fluctuation can mitigate the impact of warming on microbial communities.

Therefore, in *P. acuta*, especially in the ±5°C group, the most sensitive to the temperature changes are likely the microbes, while the host and symbiotic algae are relatively slower to respond. However in the larger fluctuation of ±7°C, the host is the most sensitive, followed by the microbes, and finally by the symbiotic algae. Although the amplitude may increase, *P. acuta* itself may adapt, and the microbial composition may also adjust to cope with such amplitude differences (Table 1).

The bacterial composition in the two corals *S. pistillata* and *P. acuta* showed different patterns (Fig. 6a). In *S. pistillata, Proteobacteria* was dominant, and *Spirochaetae* appeared, though it did not appear in *P. acuta*. In *P. acuta, Proteobacteria* and *Chlamydiae* were dominant in the composition (Fig. 6b). Although *Proteobacteria* was dominant in both *S. pistillata* and *P. acuta*, the relative abundance of *Alphaproteobacteria* and *Gammaproteobacteria* were different in the two corals. Under the control treatment, *S. pistillata* exhibited a higher relative abundance of *Gammaproteobacteria* than *Alphaproteobacteria*. However, in the ±5°C and ±7°C treatments, there was an increased relative abundance of *Alphaproteobacteria*. Under every treatment, *P. acuta* exhibited a higher relative abundance of *Gammaproteobacteria* than *Alphaproteobacteria*, though this was especially true in the ±5°C treatment.

**Fig. 6.**
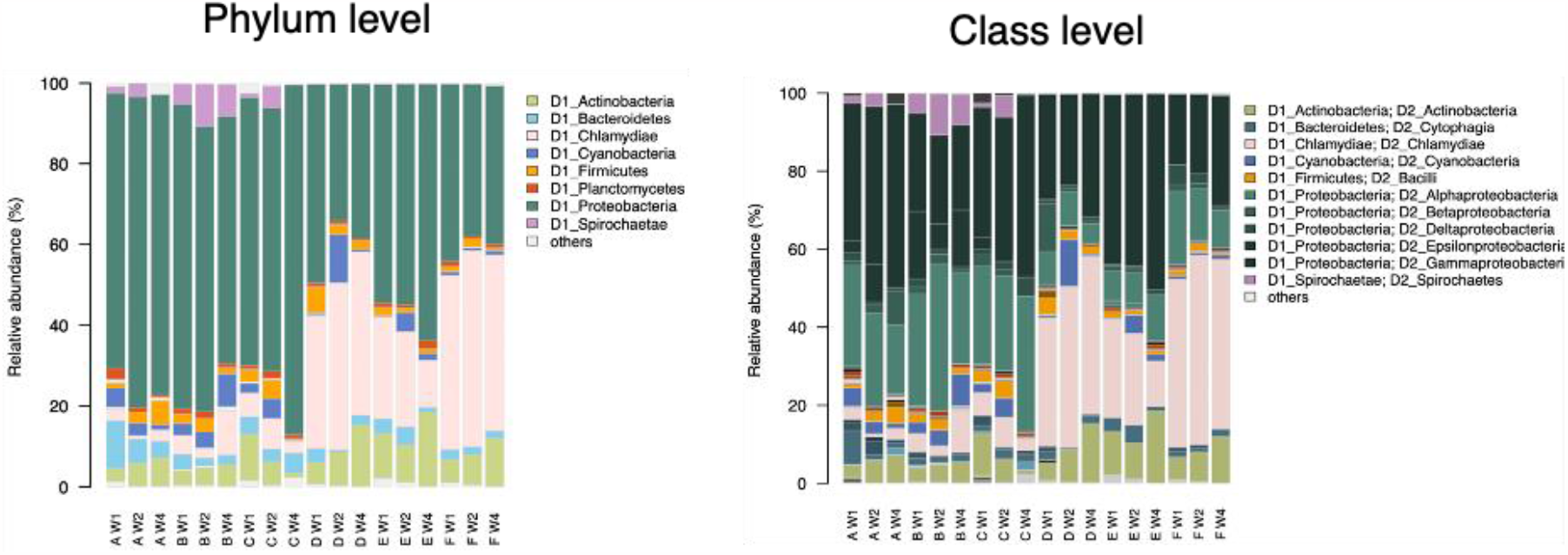
Bacterial composition at the phylum and class levels.

According to the beta diversity results, the bacterial composition in *S. pistillata* manifested differences in the second week of daily temperature fluctuation treatments (Fig. 5a). Based on the indicator bacteria results of the second week in *S. pistillata* (Fig. 7), the KTUs with significant differences among the groups were mainly dominated by the phyla *Alphaproteobacteria* and *Gammaproteobacteria*. For example, in both the ±5°C and ±7°C groups, *Rhodobacteraceae*, which belong to *Alphaproteobacteria*, were observed to have significant relative abundance differences. Additionally, in the ±7°C group, the abundances of *Acinetobacter* and *Endozoicomonas* belonging to *Gammaproteobacteria* were also significantly different.

**Fig. 7.**
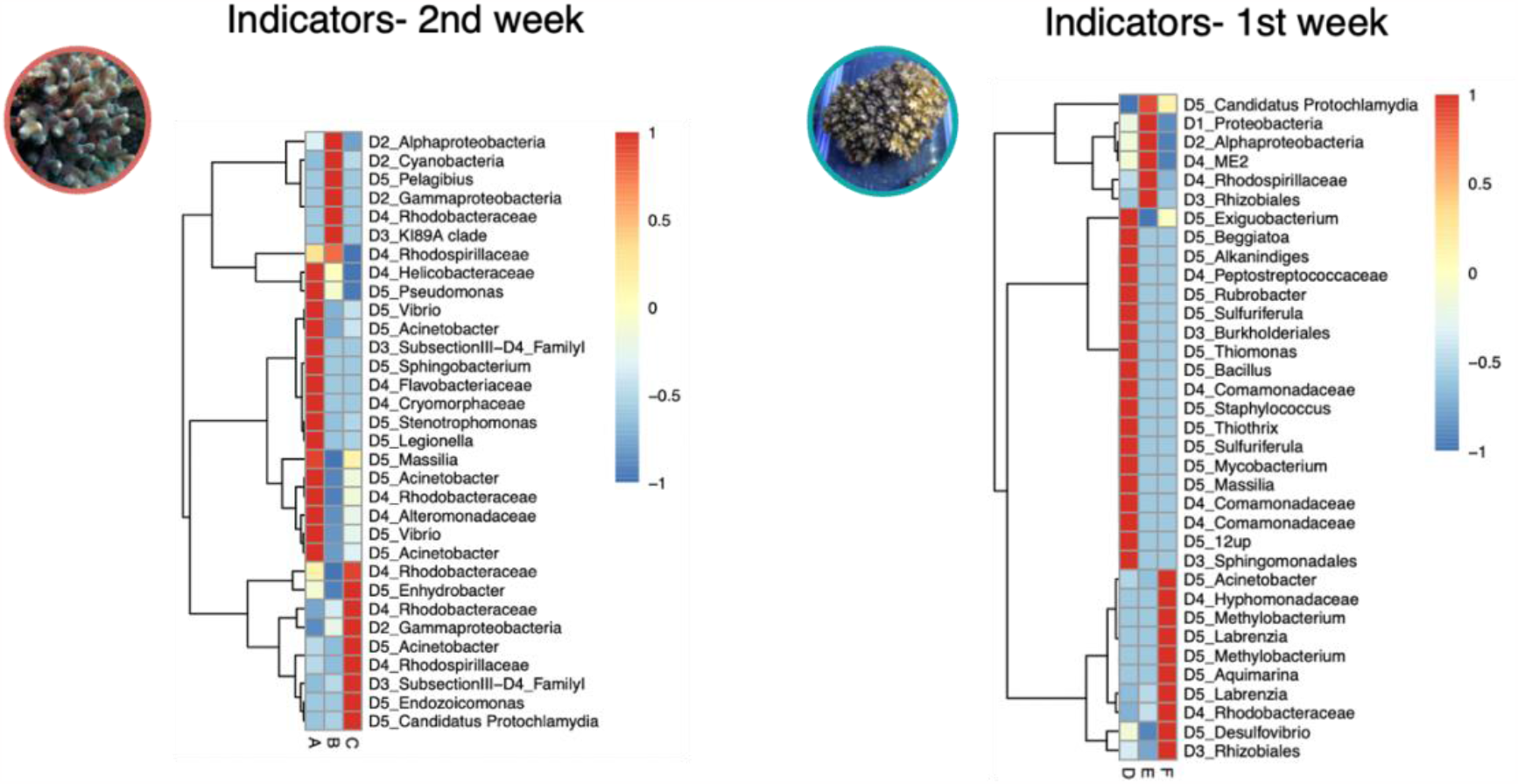
Indicator bacterial KTUs with significant differences among the groups showed in the two corals.

In *P. acuta*, the bacterial composition showed the most sensitive response after treatment, and its sensitivity to increasing temperature and diurnal temperature range variation could be detected within the first week (Fig. 5b). According to the indicator bacteria results of the first week in *P. acuta* (Fig. 7), the KTUs with significant differences among the groups were mainly dominated by the phylum *Alphaproteobacteria*. For example, in the ±5°C group, *Rhodospirillaceae* and *Rhizobiales* were observed to have significant relative abundances, and in the ±7°C group, *Hyphomonadaceae, Methylobacterium, Labrenzia*, and *Rhodobacteraceae* were observed to be significant. In addition to *Alphaproteobacteria, Acinetobacter* belonging to *Gammaproteobacteria* was also significantly different in the ±7°C group.

Interestingly, these indicator bacteria with significant differences, such as *Acinetobacter* spp., appeared more evenly distributed across the samples. Specifically, in *S. pistillata, Endozoicomonas* only appeared at a relatively high abundance in two samples of the ±7°C group, while *Acinetobacter* appeared more evenly across all samples of both the ±5°C and ±7°C groups (Fig. S4a). Additionally, there were more KTUs belonging to the genus *Acinetobacter* than *Endozoicomonas*. Similarly, in *P. acuta, Acinetobacter* appeared more evenly in all samples of the ±5°C and ±7°C groups compared to other indicator bacteria (Fig. S4b).

Comparing the core bacterial KTUs of the two hosts across different treatment groups and time points, we found that 66 genera were present in *S. pistillata* and 46 were present in *P. acuta* (Fig. S5). However, only 14 of these core KTUs were shared by both species (Table S1). Interestingly, the bacterial genera belonging to *Rhodobacteraceae* and *Acinetobacter* were among these 14 core KTUs. This result suggests that *Rhodobacteraceae* and *Acinetobacter* are relatively stable core bacteria in corals, but their relative abundance is influenced by daily temperature fluctuation, making them potential indicator bacteria for both of these coral species under heat stress.

## 4. Discussion

### 4.1 Diurnal temperature fluctuation may mitigate coral heat buildup

Coral species inhabiting the intertidal zone of coasts and estuaries have the capacity to withstand significant fluctuations in environmental conditions, particularly temperature, due to the diverse and extensive natural variations of the habitat (Somero, 2002). Although the effects of short temporal variation on a daily to weekly scale are rarely studied, Safaie et al. (2018) suggest that daily temperature fluctuations may reduce the heat accumulation on corals. Also, Oliver and Palumbi (2011) suggest that corals that experience significant fluctuations in daily temperatures may exhibit increased resistance to heat stress. Their results from *Acropora hyacinthus* tank experiments suggest that samples that experience larger diurnal temperature variations have higher heat tolerance compared to those that experience smaller diurnal temperature variations. Additionally, Pineda et al. (2013) analyzed the 2010 bleaching event that occurred in the central area of the Red Sea and found that *S. pistillata* located in the nearshore areas with large temperature fluctuations had a lower mortality rate than *S. pistillata* located in areas with smaller temperature variations.

Different coral species have varying responses and tolerances to heat stress. At the nuclear powerplant outlet shallow reef in Taiwan, Keshavmurthy et al. (2014) found that *P. acuta* populations acclimatized better than *S. pistillata* under the constant warming SST. This research also demonstrated, through the use of a tank experiment, that *P. acuta* is more tolerant to temperature fluctuations than *S. pistillata* under different daily temperature treatments. The plasticity in corals’ responses to temperature changes may result from adaptations at the genetic level within populations, and individuals may undergo acclimation through physiological changes that involve regulating gene expression or cell response (Middlebrook et al., 2008; Barshis et al., 2013).

### 4.2 Tripartite dynamic response within the holobiont to the daily temperature fluctuations

The tripartite dynamic response within the holobiont to the daily temperature fluctuations did not immediately synchronize. In other words, the response times for the host, symbiotic algae, and bacteria varied and this timing appeared to be influenced by coral species. When the daily temperature fluctuation is small, microbes may be more sensitive or as sensitive as the host to temperature change, depending on the coral species; when the fluctuation is large, the host will be more sensitive than the microbes. Regardless of species, symbiotic algae show the slowest response to temperature fluctuations.

Individual holobiont partners may respond to heat stress differently. For example, the coral host would likely respond to heat stress before the symbiotic algae. Leggat et al. (2011) examined the expression of genes involved in stress response and carbon metabolism in the coral *Acropora aspera* and its symbiotic algae under a stable temperature increase. They found that although there was no significant decline in Fv/Fm when corals were incubated in 34°C seawater for 2 days, there were significant changes in host gene expression, but very little change in the expression of metabolic genes in the symbiotic algae. In a similar study that also incorporated the bacteria of the holobiont, Li et al. (2021) used stable warming conditions to treat *Pocillopora damicornis* and observed successive changes in the host, symbiotic algae, and bacterial community. Although they did not consider diurnal temperature variation, they still observed changes in host gene expression and bacterial community that happened earlier and with more pronounced changes than that of symbiotic algae in response to warming. Genetic variations within host populations could also lead to different responses. Humanes et al. (2022) exposed an *A. digitifera* population to heat stress, and the colonies that were least tolerant perished, while the most tolerant survived. Surprisingly, this discrepancy did not seem to connect to the specific type of symbiotic algae, indicating that the coral itself possessed greater or lesser heat tolerance.

In sum, it can be inferred from all these results and the results of this study that when coral holobionts are under heat stress, there are significant differences in heat tolerance and heat resilience in different coral hosts. Although the host and its bacterial community are the first to respond, *P. acuta* can respond quickly and adjust before the symbiotic algae respond to heat stress, whereas *S. pistillata* lacks the ability to make adjustments, leading to subsequent bleaching.

### 4.3 Tank experiments without diurnal temperature fluctuation may find enlarged microbial dynamics

Previous studies have suggested that since high-frequency temperature variability reduces the heat accumulation in corals (Safaie et al., 2018), the diurnal temperature fluctuation may also reduce the impact of warming on microbial communities. However, according to this research, a smaller variation of microbial compositions was observed in both treatments with daily temperature fluctuation. These differing results may offer some guidance for future tank experiments: if stable warming is used to observe changes in microbial communities without considering diurnal temperature fluctuations, the results may increase the variations in microbial populations.

### 4.4 Different coral species may adopt different adaptation and survival strategies in response to diurnal temperature oscillations and warming

The microbiomes of coral mucus, tissue, and skeleton exhibit differences in their microbial community composition, richness, and sensitivity to host-specific and environmental factors. The microbiome can be phylosymbiotic, in which the composition and richness may also reflect the phylogenetic relationship of the coral host (Pollock et al., 2018). The results of this study showed that different coral species and their microbial components may adopt distinct adaptation and survival strategies in response to daily temperature fluctuations and warming. Additionally, it can be inferred that there may be causal relationships among hosts, microbes, and symbiotic algae in response to warming and diurnal temperature fluctuations. In species that are more sensitive to heat, changes in host physiology may affect their symbiotic microbes or algae; in more heat-tolerant species, even though the host physiology would be affected by the heat stress, the host may respond promptly, minimizing the influence on the microbes and symbiotic algae. However, this still requires further confirmation.

### 4.5 Potential indicator coral-associated bacterial species

Coral reefs are currently encountering unparalleled challenges at both local and global levels. There is an urgent requirement for easily detectable and highly sensitive indicators of ecosystem stress to support the development of efficient management and restoration techniques. Many studies have investigated microbial indicators for environmental perturbations in reef ecosystems or corals (Ziegler et al., 2017; Glasl et al., 2019)

Both bacteria *Rhodobacteraceae* and *Moraxellaceae* are often found in coral tissues and mucus (Pollock et al., 2018; Ostria-Hernández et al., 2022). Kuek et al. (2022) found that *Rhodobacteraceae* was one of the dominant bacteria families in the mucus and tissue of both *P. acuta* and *S. pistillata*. Grottoli et al. (2018) subjected both *Acropora millepora* and *Turbinaria reniformis* to heat and acidification treatments, and they found that although *Rhodobacteraceae, Acinetobacter* (*Moraxellaceae*), and *Coxiella* bacteria were present in corals under both control and treatment conditions, their relative abundance changed significantly due to the treatments. In Ziegler et al. (2019), a transplant experiment involving *Acropora hemprichii* and *Pocillopora verrucosa* was conducted in three locations which were experiencing different levels of anthropogenic disturbance.They found that after exposing the corals to significant levels of anthropogenic disturbance, the relative abundance of *Rhodobacteraceae* and *Moraxellaceae* in their microbiomes would increase. Li et al. (2021) also found that, in *Pocillopora damicornis*, as the temperature increased, the relative abundance of *Rhodobacteraceae* increased while *Endozoicomonas* decreased. These results are similar to the bacterial community changes observed in our experiment on *P. acuta*. Therefore, we can conclude that bacterial families such as *Rhodobacteraceae* and *Moraxellaceae*, including *Acinetobacter*, are commonly present in corals, and their responses to environmental changes can be rapid and prominent, making them potential indicators of whether corals are experiencing heat stress.

The genus *Acinetobacter*, belonging to *Moraxellaceae*, is reported to be one of the dominant bacterial genera in multiple coral species from many regions (Carlos et al., 2013; Littman et al., 2009; Li et al., 2014; McKew et al., 2012; Morrow et al., 2012,). It may act as a frontline defense and protect the coral holobiont against pathogens that are resistant to multiple antibiotics (Shnit-Orland and Kushmaro, 2009; Yang et al., 2017), but it is also regarded as a potential pathogen to corals (Sweet et al., 2013). Besides this, *Acinetobacter* may also interact with metabolic compounds produced by coral hosts (Ding et al., 2016; Horinouchi et al., 1997). Inprevious studies, *Acinetobacter* has been found to have a broader geographic range and less host specificity than the coral symbiotic bacteria *Endozoicomonas* (Yang et al., 2020, 2017). In this study, *Acinetobacter* is represented as one of the core coral bacterial genera in both *S. pistillata* and *P. acuta* in all temperature fluctuation treatments, indicating that it could be more flexible than other bacteria with different coral species and temperature conditions. Additionally, compared to other potential indicator bacteria, such as *Rhodobacteraceae, Endozoicomonas*, and *Vibrio*, the relative abundance of *Acinetobacter* was well-distributed in most of the samples, rather than only being found in one or two samples of a treatment. Therefore, based on this result, we suggest that *Acinetobacter* is a potential bacterial indicator for coral hosts under large daily temperature fluctuations with increasing temperature.

Although the effect of diurnal temperature oscillations on reducing heat accumulation in corals still needs further study, the current experiment indicates that the response rates of coral hosts, symbiotic algae, and microbiomes vary depending on temperature oscillation ranges and coral species. We also found that *Acinetobacter*, which is a *Gammaproteobacteria*, may be more suitable as a microbial indicator for daily temperature fluctuation in coral holobionts than the common coral symbiotic bacteria *Endozoicomonas*. This study sheds light on the causal relationships among the host, symbiotic algae, and microbes in corals under extreme climate and warming pressures. Hence, we believe that these findings can also facilitate future assessments of the potential application of probiotics for coral restoration, as well as the use of indicator microbes for monitoring purposes.

## Acknowledgments

We thank Dr. Sen-Lin Tang of Academia Sinica for the equipment support. We also thank Mr. Justin Pelofsky of Third Draft Editing for his English language editing.

## Contribution

Yunli Eric Hsieh: performed molecular experiment, data analysis, sequences submission, manuscript writing

Chih-Ying Lu: performed all the tank experiments and physiological experiments

Po-Yu Liu: Data analysis and statistical Analysis

Sung-Yin Yang: manuscript writing

Chia-Min Kao: molecular experiment, data analysis

Chien-Yi Wu and Jing-Wen Michelle Wong: assisting with tank experiment and physiological experiment

Shinya Shikina, Tung-Yung Fan: sampling and preceding operation for tank experiments

Shan-Hua Yang: conceived of the idea, designed research, manuscript writing

## Data availability

The original data set presented in the study is publicly available. These data can be found at NCBI under BioProject accession number: PRJNA949725.

