## Supplementary material for "Successive responses of three coral holobiont components (coral hosts, symbiotic algae, and bacteria) to daily temperature fluctuations"

**Figures:**

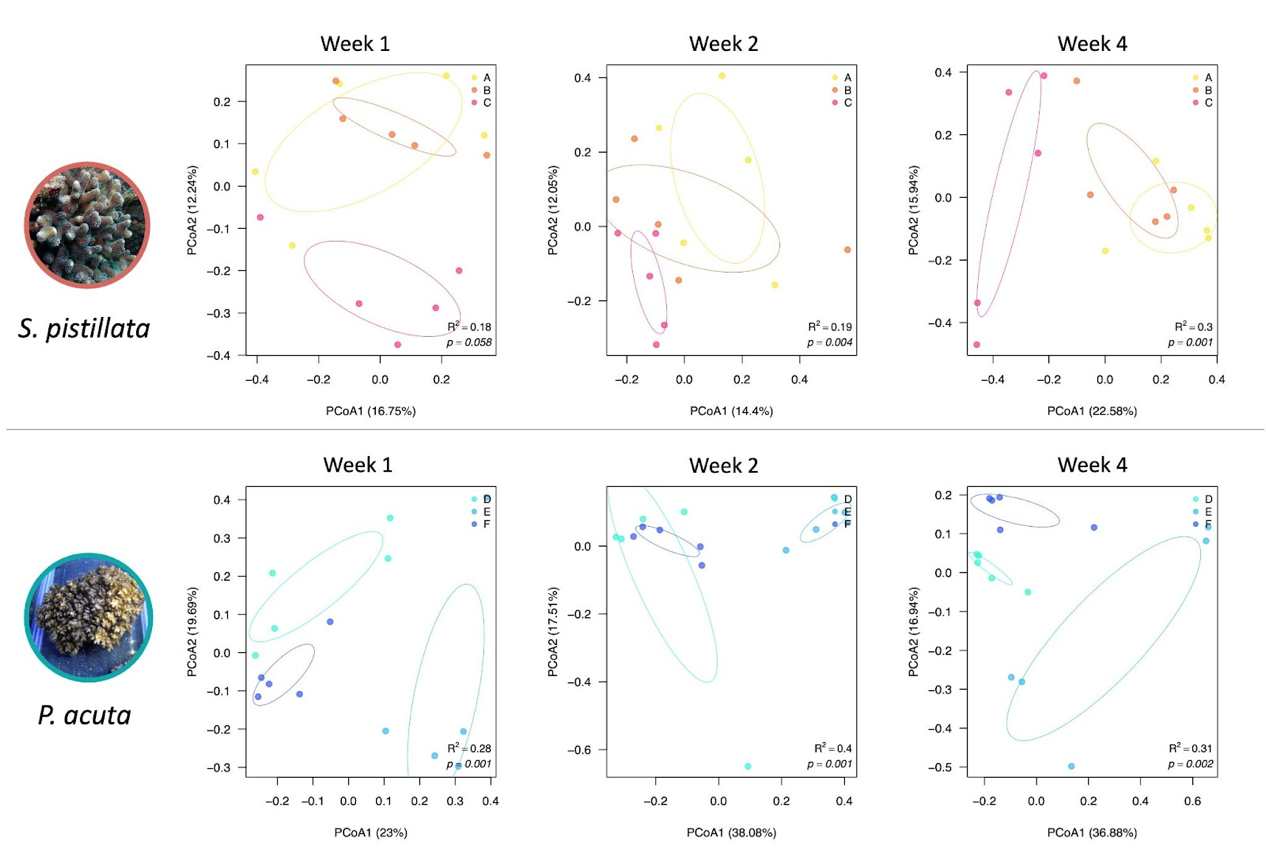

**Fig. S1.** Bacterial dynamics in the two corals at week 1, week 2 and week 4. *S. pistillata* is in groups A, B, and C; and *P. acuta* is in groups D, E, and F. Groups A and D are the treatment of 26-29°C, groups B and E are the treatment of 26±5°C-29±5°C, and groups C and F are the treatment of 26±7°C-29±7°C.

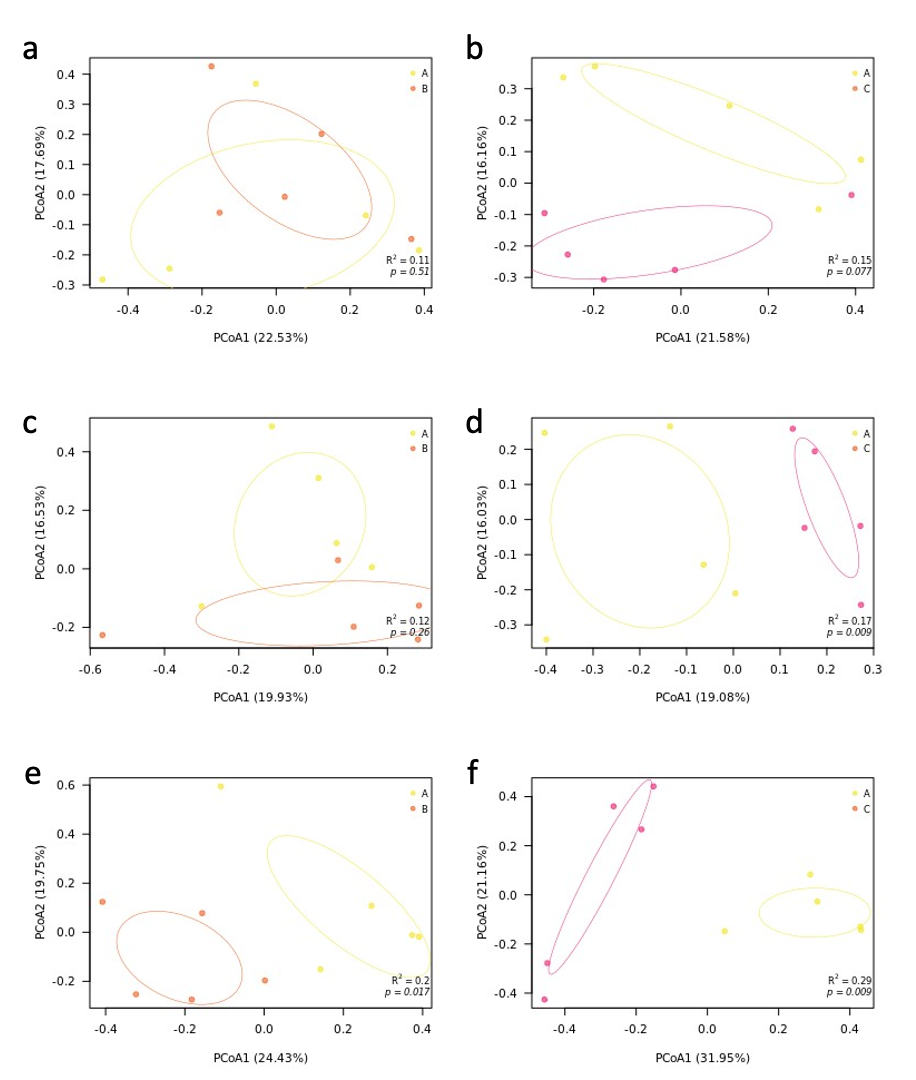

**Fig. S2.** Comparison of *S. pistillata* associated bacterial composition between control and fluctuation treatments. Panels a, c, e show ±5°C group to control in week 1, week 2 and week 4, respectively; Panels b, d, f show ±7°C group to control in week 1, week 2 and week 4, respectively.

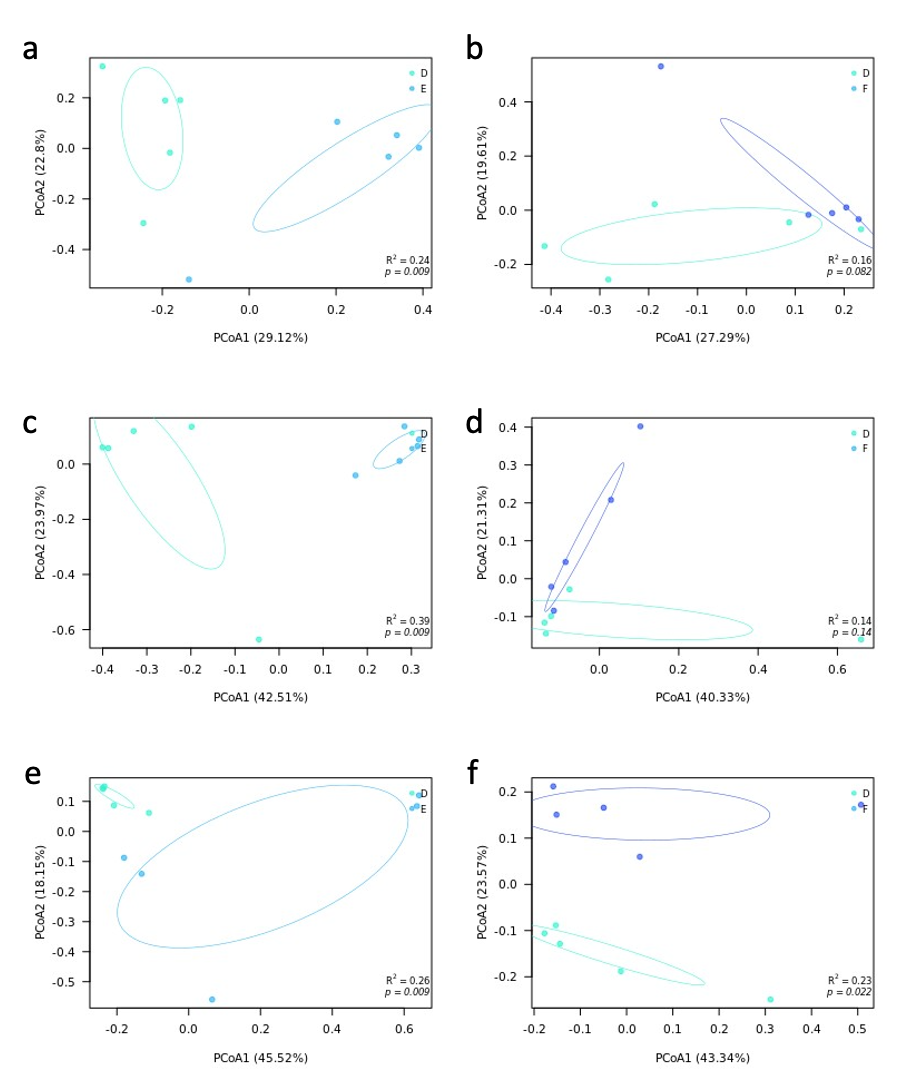

**Fig. S3.** Comparison of *P. acuta* associated bacterial composition between control and fluctuation treatments. Panels a, c, e show ±5°C group to control in week 1, week 2 and week 4, respectively; Panels b, d, f show ±7°C group to control in week 1, week 2 and week 4, respectively.

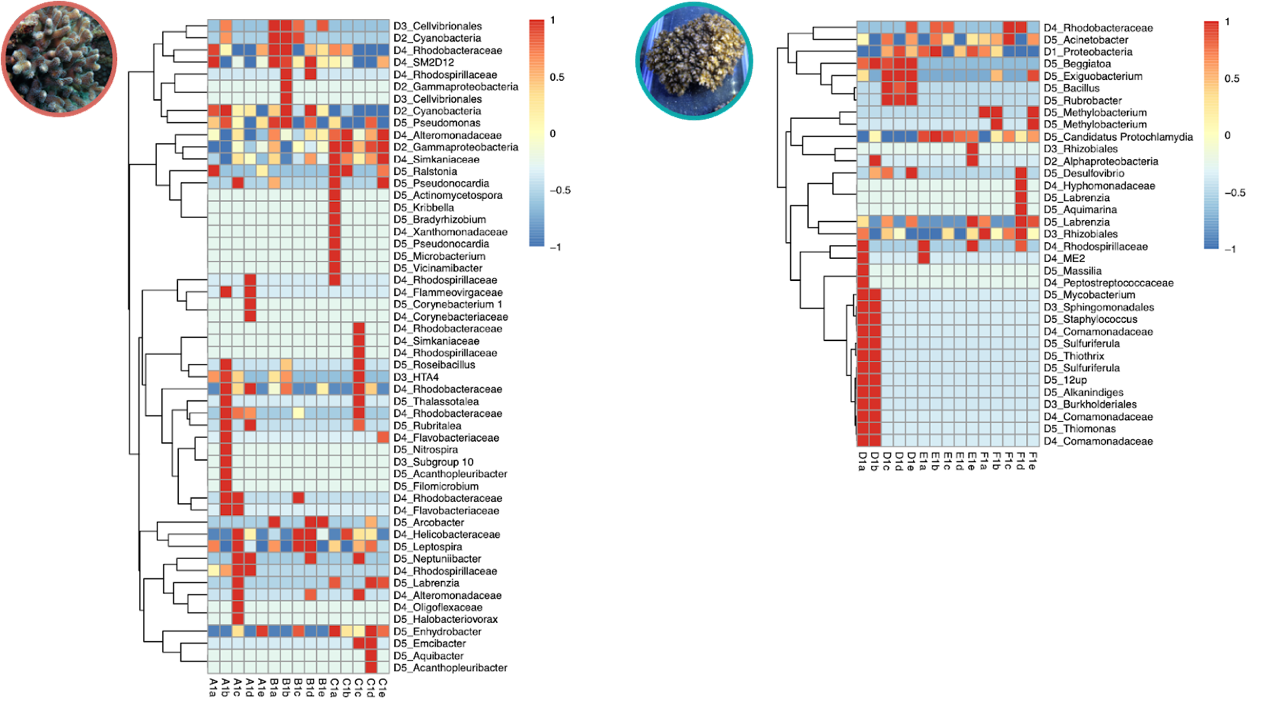

**Fig. S4a** Indicator bacteria represented in each sample of two corals in the first week. Colors indicate standardized bacterial relative abundance.

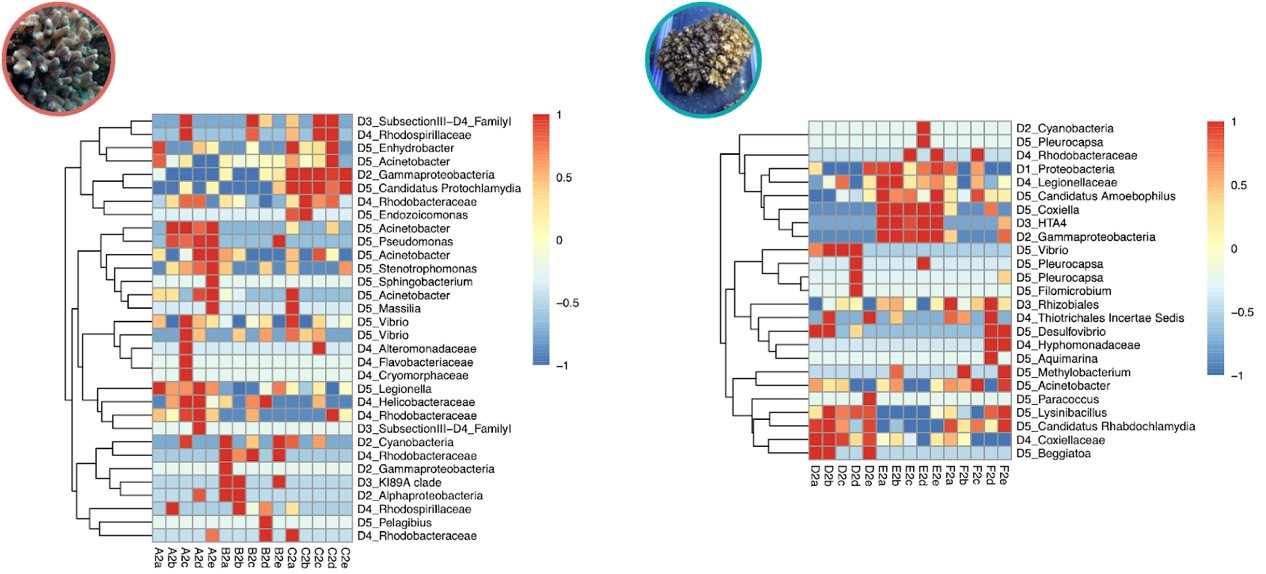

**Fig. S4b.** Indicator bacteria represented in each sample of two corals in the second week. Colors indicate standardized bacterial relative abundance.

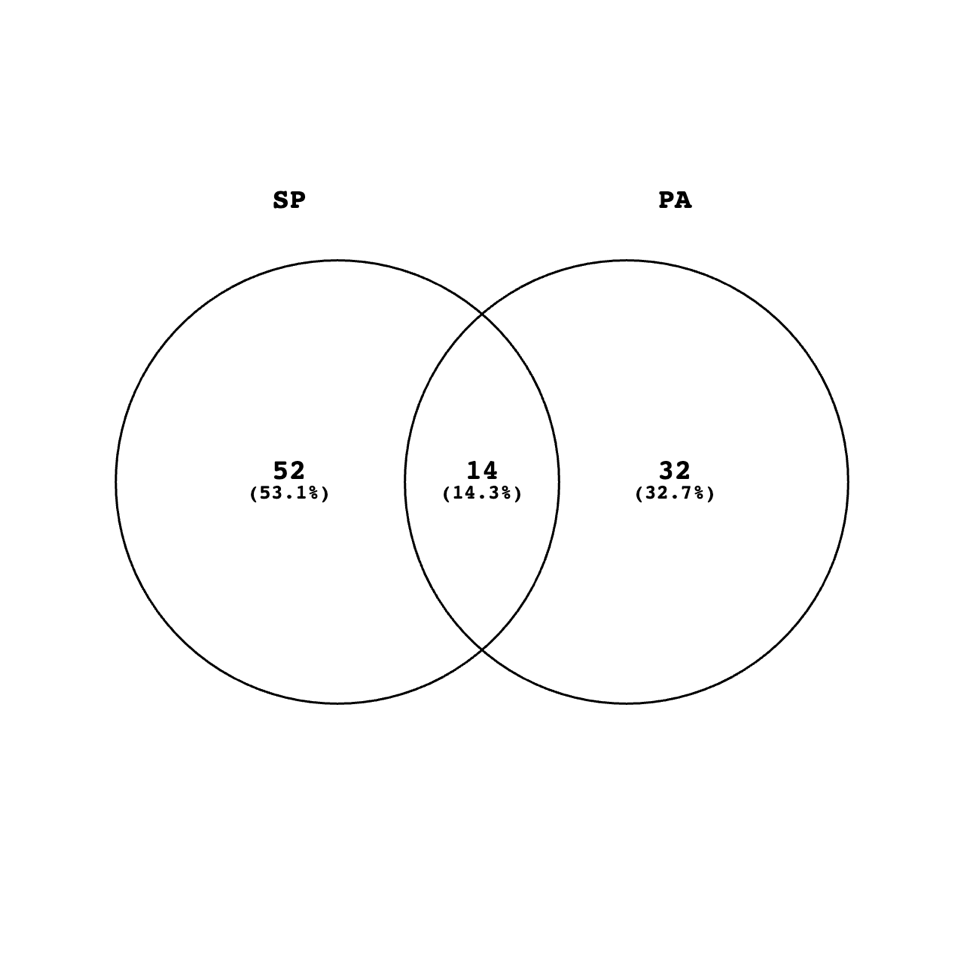

**Fig. S5.** Number of the core bacterial KTUs in both corals *S. pistillata* (SP) and *P. acuta* (PA).

**Table:**

**Table S1.** Taxa of the 14 core KTUs in both corals *S. pistillata* (SP) and *P. acuta* (PA).

| KTU | Phylum | Class | Order | Family | Genus |
| --- | --- | --- | --- | --- | --- |
| a7f393e632bf5472cbb086f93db2be65 | TM6 | TM6 | TM6 | TM6 | TM6 |
| 69c02a17ec051dacffd71a681c5ebf96 | *Proteobacteria* | *Alphaproteobacteria* | *Rhizobiales* | *-* | *-* |
| 45683537c78751927d91c137b869783f | *Proteobacteria* | *Alphaproteobacteria* | *Rhizobiales* | *Hyphomicrobiaceae* | *Filomicrobium* |
| 310e99bf8fffc716dcf3358f9a7a6b0c | *Proteobacteria* | *Alphaproteobacteria* | *Rhodobacterales* | *Rhodobacteraceae* | *Labrenzia* |
| 129314615388e9f986a235cf34829410 | *Proteobacteria* | *Betaproteobacteria* | *Burkholderiales* | *Comamonadaceae* | *Delftia* |
| 114beb9ade7b6e9f2b6476e765084c0f | *Proteobacteria* | *Gammaproteobacteria* | *-* | *-* | *-* |
| 4f3b636d4dbb1e7b7968e5892a6316d3 | *Proteobacteria* | *Gammaproteobacteria* | *Pseudomonadales* | *Moraxellaceae* | *Acinetobacter* |
| ffc20b49d9596ca6861a06bc5ddd96a5 | *Proteobacteria* | *Gammaproteobacteria* | *Legionellales* | *Coxiellaceae* | *Coxiella* |
| d2535f4db5b2b496347303b1facfbf8c | *Proteobacteria* | *Gammaproteobacteria* | *Legionellales* | *Coxiellaceae* | *Coxiella* |
| 66567bcb25aba2c5fa20874b7c002797 | *Actinobacteria* | *Actinobacteria* | *Propionibacteriales* | *Propionibacteriaceae* | *Propionibacterium* |
| 09f995675b2073cece4f46980fbd3370 | *Actinobacteria* | *Actinobacteria* | *Micrococcales* | *Micrococcaceae* | *Rothia* |
| 8f0f040f00b3adbd38e00a781fd70a09 | *Chlamydiae* | *Chlamydiae* | *Chlamydiales* | *Simkaniaceae* | *-* |
| e03a3ce0fcae1838fd6a5684ab9c2aed | *Chlamydiae* | *Chlamydiae* | *Chlamydiales* | *Simkaniaceae* | *Simkania* |
| 8d694120e37d4485b0cb68525380ddcd | *Chlamydiae* | *Chlamydiae* | *Chlamydiales* | *Simkaniaceae* | Candidatus *Rhabdochlamydia* |
